# Ablation of *Arf*-expressing cells ameliorates cigarette smoke-induced lung dysfunction in mice

**DOI:** 10.1101/809160

**Authors:** Ryuta Mikawa, Tadashi Sato, Yohei Suzuki, Hario Baskoro, Masataka Sugimoto

**Affiliations:** Research Institute, National Center for Geriatrics and Gerontology; Department of Pulmonary Medicine, Juntendo University Graduate School of Medicine; Department of Molecular Aging Research, Nagoya University Graduate School of Medicine

## Abstract

Senescent cells accumulate in tissues during aging or pathological settings. The semi-genetic or pharmacological targeting of senescent cells revealed that cellular senescence underlies many aspects of the aging-associated phenotype and diseases. We previously reported that cellular senescence contributes to aging- and disease-associated pulmonary dysfunction. We herein report that the elimination of *Arf*-expressing senescent cells ameliorates cigarette smoke-induced lung pathologies in mice. Cigarette smoke induced the expression of *Ink4a* and *Arf* in lung tissue with concomitant increases in lung tissue compliance and alveolar airspace. The elimination of *Arf*-expressing senescent cells prior to cigarette smoke exposure protected against these changes. Furthermore, the administration of cigarette smoke extract lead to pulmonary dysfunction, which was ameliorated by subsequent senescent cell elimination. Collectively, these results suggest that senescent cells are a potential therapeutic target for cigarette smoking-associated lung disease.

## Introduction

Chronic obstructive pulmonary disease (COPD) is a severe and incurable disease with a relatively high prevalence and is currently one of the leading causes of death worldwide [1]. A major component of COPD is pulmonary emphysema, which is characterized by enlarged alveolar airspaces and alveolar wall collapse. Cigarette smoke (CS) is the dominant cause of the disease, and aging also increases the risk of COPD [2]. COPD is associated with the infiltration of inflammatory cells, which is considered to cause the accumulation of proteinases and lead to alveolar destruction [3, 4]. The inhibition of macrophage or neutrophil elastase was previously shown to be sufficient to suppress the pathologies of CS-induced emphysema in mice [5, 6]. Cellular senescence is also accelerated in emphysema patients [7]. However, the roles of cellular senescence in this disease have yet to be characterized.

Cellular senescence acts as a potent tumor suppression mechanism in mammals by halting the proliferation of cells at risk of malignant transformation [8]. Cellular senescence is mediated by the *Cdkn2a* locus, which encodes two important tumor suppressors, namely, p16^Ink4a^ and p19^Arf^ (p14^ARF^ in humans) [9, 10]. These proteins indirectly activate the Rb and p53 tumor suppressors, respectively, thereby inducing and maintaining cell cycle arrest during cellular senescence. Besides permanent cell cycle arrest, it has become apparent that senescent cells produce a series of cytokines, chemokines, and extracellular proteinases, thereby non-cell autonomously affecting the functions of their surrounding non-senescent cells. These are collectively called SASP (the senescence-associated secretory phenotype), and increasing evidence suggests that SASP mediates the deleterious effects of cellular senescence and is involved in a number of pathologies in both humans and mice [11].

Senescent cells accumulate in several tissues during aging [12, 13], and are considered to contribute to aging-associated pathologies. Recent studies on senolysis (senescent cell elimination) using semi-genetic and pharmacological approaches revealed that the elimination of senescent cells from tissues ameliorates a number of aging-associated tissue dysfunctions and diseases [11, 14] and even extends the lifespan of mice [15]. Cellular senescence is also involved in lung disease, and bleomycin-induced idiopathic pulmonary fibrosis (IPF) was shown to be alleviated by the elimination of senescent cells in INK-ATTAC mice or the administration of the senolytic drugs dasatinib and quercetin [16]; however, the efficacy of these drugs in human IPF patients currently remains unclear [17].

We also established transgenic mice (ARF-DTR mice) that express luciferase and diphtheria toxin (DT) receptors under the control of the *Arf* promoter/enhancer, thereby enabling the detection and ablation of *Arf*-expressing cells by *in vivo* imaging and DT, respectively [18]. Using ARF-DTR mice, we showed that the elimination of *Arf*-expressing cells restored pulmonary function in aged animals [19]. Furthermore, the pre-elimination of *Arf*-expressing cells by DT protected lung tissue against elastase-induced lung injury [20]. This effect may be mediated by reduced pulmonary inflammation exemplified by an increased number of alveolar macrophages. Similar findings were also obtained using the anti-apoptotic Bcl-2 family protein inhibitor ABT-263, which has been shown to exhibit senolytic activity in mice [21]. Collectively, these findings suggest that the elimination of senescent cells exerts protective effects against elastase-induced pulmonary emphysema. However, it currently remains unclear whether the elimination of senescent cells is also effective in CS-induced pulmonary injury, which is more closely related to common human diseases.

In the present study, we investigated the effects of *Arf*-expressing cells on CS-induced lung pathologies in mice. Chronic CS exposure resulted in pulmonary dysfunction and morphological abnormalities, and these CS-induced phenotypes were mitigated in mice in which *Arf*-expressing cells were eliminated prior to CS exposure. We also showed that the elimination of *Arf*-expressing cells following lung injury was beneficial in cigarette smoke extract (CSE)-administered mice. Collectively, these results imply that senolysis has potential as a therapeutic approach for human emphysema.

## Materials and methods

### Animals

All animal experiments were approved by and conducted in accordance with guidelines established by the National Center for Geriatrics and Gerontology Animal Ethics Committee. ARF-DTR mice [19] and littermate mice (wild-type) with the C57BL/6J background were maintained under specific pathogen-free conditions with a 12-hour light/dark cycle, constant temperature, and *ad libitum* access to food (CE-2; CLEA Japan) and water. DT was intraperitoneally injected at a concentration of 50 μg/kg body weight.

### CS and CSE treatment

CS was prepared using commercially marketed Peace non-filter cigarettes (Japan Tobacco) containing 29 mg of tar and 2.5 mg of nicotine per cigarette. CS exposure was performed on a tobacco smoke inhalation experiment system for small animals (Sibata Scientific Technology) [22, 23]. CS was diluted with compressed air to 3.5 % and exposed to mice at a rate of 6 puffs per minute with a 15-mL stroke volume. Mice were exposed to diluted CS or fresh air (control group) for 30 minutes per day, 5 days per week for 4 weeks.

CSE was prepared according to previously described methods [24, 25]. Smoke from 8 Peace non-filter cigarettes was drawn through 100-mL sterile phosphate-buffered saline (PBS) using a peristaltic pump at a rate of 1.0 mL per minute. Each cigarette was burned for approximately 6 minutes. CSE was cleared by centrifugation and 50 μL was intranasally administered to mice. CSE was freshly prepared prior to each set of experiments.

### *In vivo* imaging analysis

An *in vivo* luciferase imaging analysis was performed using the IVIS imaging system (Perkin Elmer). Mice were ventrally shaved and anesthetized with isoflurane (Wako Pure Chemicals Industries), and this was followed by an intraperitoneal injection of 150 mg/kg body weight of luciferin (VivoGlo; Promega). Luciferase activity was monitored 10 minutes after the luciferin injection.

### Morphometric analysis

All histopathological analyses were performed in a blinded manner. Lungs were fixed with Mildform®20N (Wako Pure Chemicals Industries) at 25 cmH_2_O. Paraffin-embedded tissues were sectioned (thickness of 5 μm) and stained with hematoxylin and eosin. At least eight randomly selected fields per mouse were photographed, and mean linear intercepts were calculated. In the measurement of mean linear intercepts, test lines were randomly drawn on images, and the length between two sequential intersections of the alveolar surface with the test line was measured. Airway and vascular structures were eliminated from the analysis.

### RNA analysis

A real-time PCR analysis was performed as previously described [26]. Total RNA was isolated from lung tissues using the PureLink® RNA Mini kit (Thermo Fischer Scientific), and reverse-transcribed using the PrimeScript RT reagent kit with a gDNA eraser (TAKARA BIO) according to the manufacturer’s instructions. PCR was performed on a CFX Real Time System (BioRad) using the KOD SYBR qPCR mix (TOYOBO). The following primers were utilized for the amplification of specific genes: *Arf*; sense 5’-GCCGCACCGGAATCCT-3’ and antisense 5’-TTGAGCAGAAGAGCTGCTACGT-3’, *Ink4a*; sense 5’-CCCAACGCCCCGAACT-3’ and antisense 5’-GCAGAAGAGCTGCTACGTGAA-3’, *Mmp-12*; 5’-CTGCTCCCATGAATGACAGTG-3’ and antisense 5’-AGTTGCTTCTAGCCCAAAGAAC-3’, *Elane*; sense 5’-CCTTGGCAGACTATCCAGCC-3’ and antisense 5’-GACATGACGAAGTTCCTGGCA-3’, *Gapdh*; 5’-AATGGTGAAGGTCGGTGTG-3’ and antisense 5’-GAAGATGGTGATGGGCTTCC-3’.

### Pulmonary function test

Pulmonary function tests were performed on a FlexiVent system (Scireq) as previously described [19, 27]. Mice were euthanized by an intraperitoneal injection of pentobarbital sodium (100 mg/kg of body weight), and connected to the FlexiVent system after tracheotomy. Mice were ventilated at a respiratory rate of 150 breaths/min with a tidal volume of 10 mL/kg against a positive end expiratory pressure of 3 cmH_2_O. Deep inflation, Snapshot-150, Quickprime-3, and a pressure-volume loop with a constantly increasing pressure were consecutively performed three times in each mouse. Static lung compliance (Cst) values were calculated by fitting the Salazar-Knowles equation to the pressure-volume loop. Dynamic lung compliance (Crs) and resistance (Rrs) values were calculated using a single frequency forced oscillation technique. Pressure, flow, and volume signals obtained from the response to a sinusoidal waveform were fit to the single compartment model using a linear regression. Tissue elastance (H) and damping (G) were obtained from respiratory system impedance data using a constant phase model. All parameters were calculated using FlexiVent software.

### Bronchoalveolar lavage fluid (BALF) analysis

BALF cells were analyzed as previously reported [28]. Briefly, BALF was prepared with 1 mL of PBS containing 5 mM EDTA, and cells were collected from BALF by mild centrifugation. Collected cells were attached to glass slides using StatSpin Cytofuge (Beckman Coulter), and subjected to modified Giemsa staining using the Diff-Quick stain kit (Sysmex).

### Statistical analysis

A one-way ANOVA was performed for comparisons of more than 2 sets of data. When the statistical model was proven to be significant, differences between combinations of the 2 groups were analyzed using a Tukey-Kramer test. A two-tailed unpaired Student’s *t*-test was used for comparisons of two sets of experimental data. Significance was represented by asterisks corresponding to \**P*<0.05, \*\**P*<0.01, and \*\*\**P*<0.001. No blinding was performed, except for histological quantifications. No statistical method was used to select the sample size.

## Results

### Cigarette smoking induces *Arf* and *Ink4a* expression in mice

To investigate the roles of p19^Arf^-expressing cells in the CS-induced lung pathology, 4-month-old ARF-DTR or wild-type mice were exposed to CS as previously described [29] for 4 weeks (30 minutes per day, 5 days per week), as shown in Fig 1A. DT was administered twice with a 2-week interval, and initial administration was performed prior to the CS exposure. We initially performed an *in vivo* imaging analysis to monitor lung luciferase expression, which was controlled by *Arf* promoter/enhancer activity in ARF-DTR mice [19]. Luciferase activity in the chest region, which represents the expression of luciferase in lung tissue [19], was slightly increased in CS-exposed ARF-DTR mice (Fig 1B). RNA was then extracted from lung tissue and the expression of *Arf* and *Ink4a* mRNA was analyzed. CS exposure resulted in an increase in both *Arf* and *Ink4a* mRNA levels (Fig 1C and 1D), which is in contrast with previous findings obtained from an elastase-induced emphysema model showing no significant change in the mRNA levels of these genes [20].

**Fig 1.**
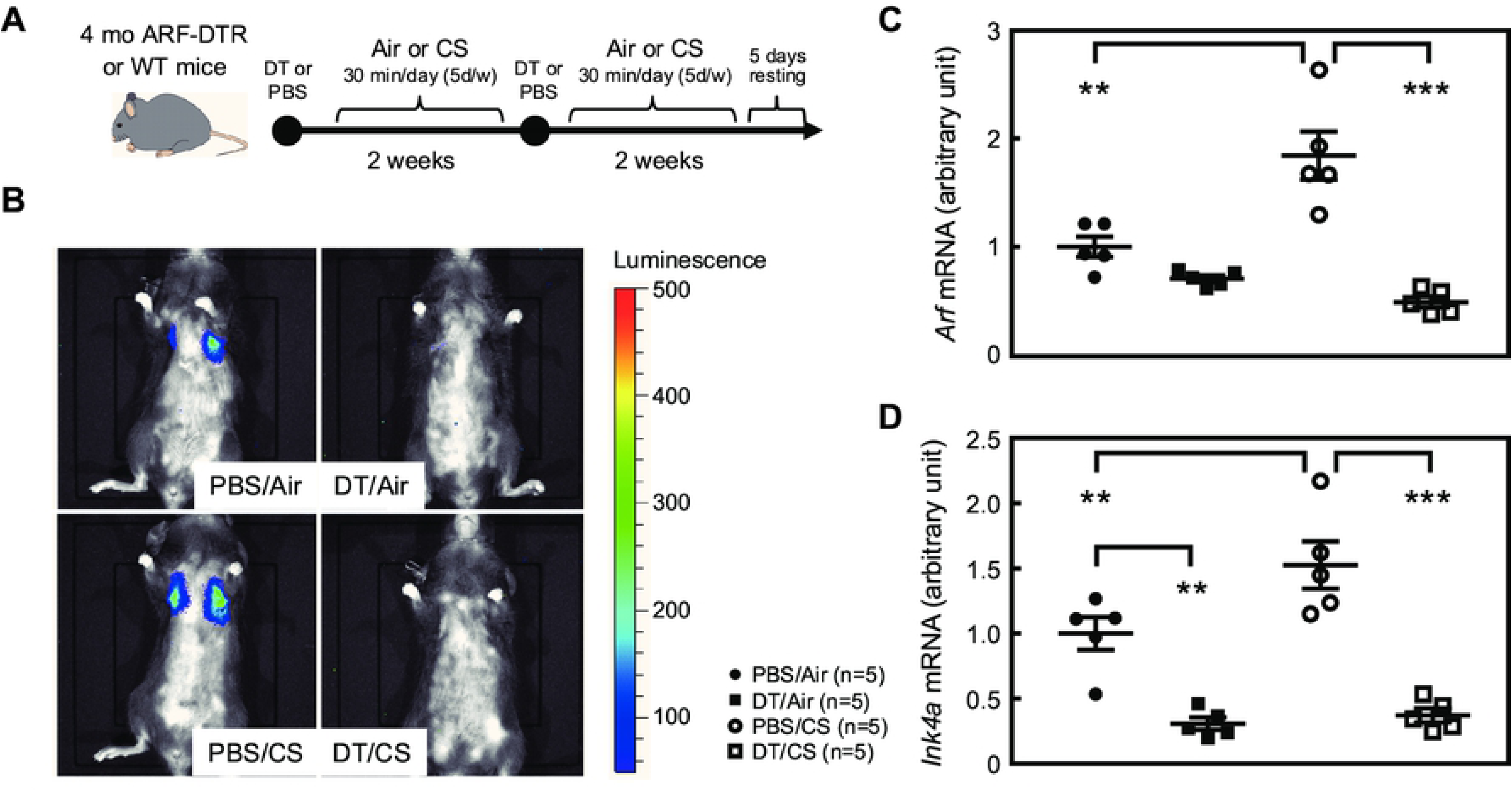
CS increased *Arf/Ink4a* levels in ARF-DTR mice. (**A**) Experimental schedule. Four-month-old ARF-DTR mice were exposed to CS for 4 weeks. DT or PBS was intraperitoneally administered twice with a 2-week interval. (**B**) An *in vivo* imaging analysis was performed 4 weeks after the CS exposure. Representative images are shown. (**C** and **D**) The expression of *Arf* (C) and *Ink4a* (D) mRNA was analyzed by real-time PCR in the lung tissue of ARF-DTR mice. Data were normalized to *Gapdh* in each group. Bars represent means ± SEM. Data were analyzed by a one-way ANOVA and Tukey post-hoc analysis. \*\**P*<0.01 and \*\*\**P*<0.001.

The administration of DT resulted in significant reductions in both *Arf* and *Ink4a* mRNA levels (Fig 1C and 1D). Consistently, luciferase activities were under the detection limit after the DT treatment (Fig 1B). On the other hand, DT had no effect on *Arf* or *Ink4a* mRNA levels in wild-type mice, which confirmed that the expression of these genes was ablated through the transgene (S1 Fig).

### Ablation of *Arf* expression attenuates CS-induced lung pathologies

We analyzed the effects of CS and DT in ARF-DTR mouse lung tissue. After a 4-week exposure to CS, lung tissues were inflated with fixative solution at a constant pressure prior to sectioning. As shown in Fig 2A, partial alveolar wall destruction was observed in the CS-exposed (PBS/CS) ARF-DTR mouse lung, which appeared to be suppressed in DT-treated ARF-DTR mice. To obtain more quantitative data, we evaluated the alveolar size by measuring the mean linear intercept (chord) length. DT did not affect the alveolar size at this age, while it was reduced by DT in the older animals [19] (Fig 2B). CS exposure resulted in a significant increase in the alveolar size, and the administration of DT suppressed this effect in ARF-DTR, but not in wild-type mice, suggesting that the elimination of *Arf*-expressing cells protected the tissue from CS-induced alveolar collapse. We previously suggested that the elimination of p19^Arf^-expressing cells suppressed elastase-induced alveolar inflammation, thereby protecting tissue from emphysema-associated pathologies [20]. This may also be the case in the CS model because the number of macrophages, which was increased in CS-exposed mice, was maintained in DT-treated animals (Fig 2C). *Mmp* (*matrix metalloproteinase*)*-12* was consistently down-regulated in lung tissue after the DT treatment (Fig 2D). DT did not markedly affect alveolar sizes, macrophage numbers, or *Mmp-12* expression in CS-exposed wild-type mice (S2 Fig). Collectively, these results suggest that similar to the elastase-induced lung injury model, the CS-induced lung pathology was attenuated by the elimination of p19^Arf^-expressing cells.

**Fig 2.**
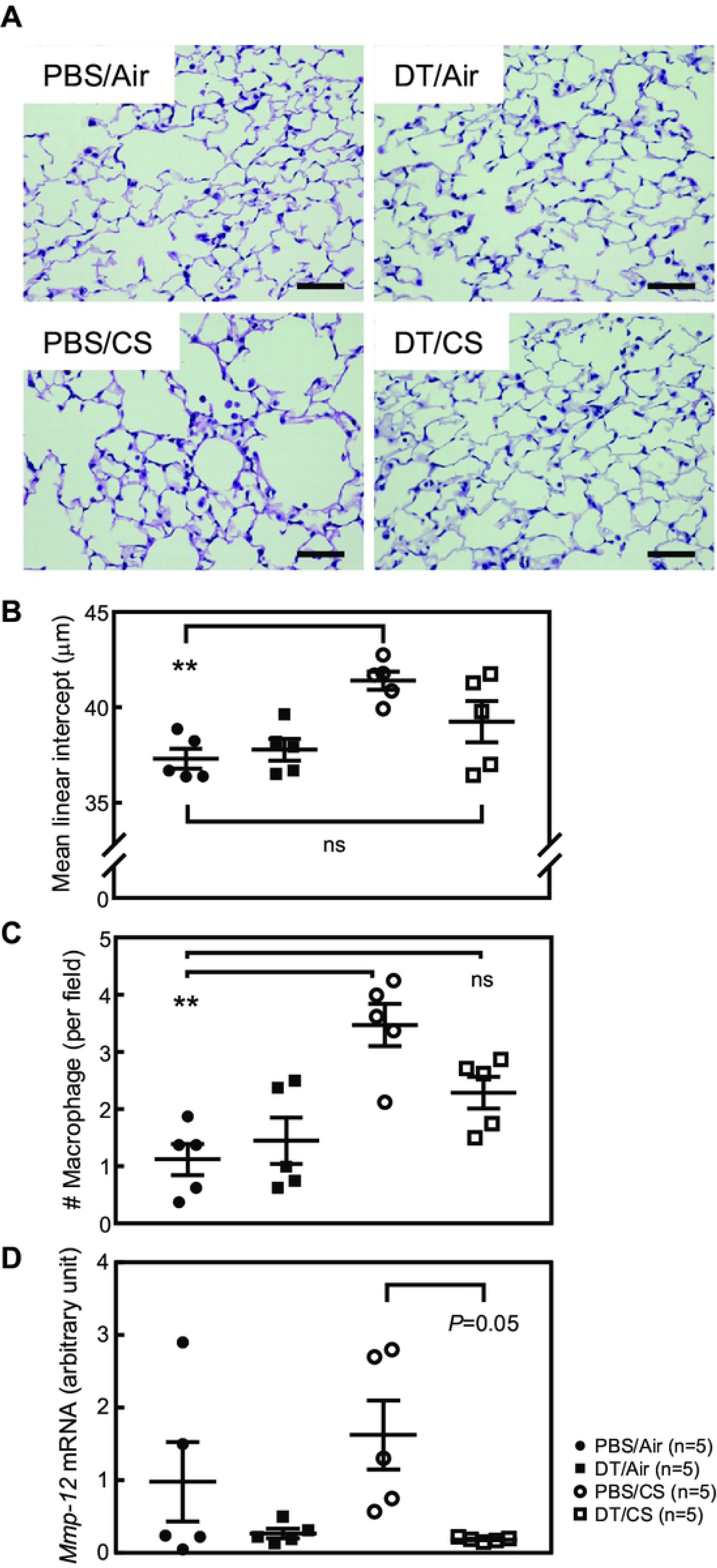
Ablation of *Arf*-expressing cells ameliorated CS-induced lung pathologies. (**A**) Representative images of ARF-DTR lung sections stained with hematoxylin and eosin. Bar; 50 μm. (**B**) Alveolar mean linear intercepts were measured. (**C**) The number of macrophages per field (×40) was counted. At least 8 fields were analyzed in each mouse. (**D**) The expression of *Mmp-12* was analyzed by real-time PCR. Data were normalized to *Gapdh* in each group. Bars represent means ± SEM. Data were analyzed by a one-way ANOVA and Tukey post-hoc analysis. \*\**P*<0.01 and ns; not significant.

### *Arf*-expressing cells facilitates CS-induced pulmonary dysfunction

The destruction of alveolar walls by CS or elastase leads to a decline in pulmonary function, which is exemplified by increased lung tissue compliance. Hence, we tested whether the elimination of p19^Arf^-expressing cells restores pulmonary function in CS-exposed ARF-DTR mice. Inspiratory capacity (IC) was significantly increased in CS-exposed mice (Fig 3A). Consistently, lung tissue compliance (Cst; static lung compliance, Crs; respiratory system compliance) was significantly increased in CS-exposed mice (Fig 3B-3D). G (tissue damping) was decreased in CS-exposed ARF-DTR mice (Fig 3G). However, the effects of CS on these parameters were diminished or prevented when DT was administered prior to the CS exposure in ARF-DTR, but not in wild-type animals (Fig 3 and S3 Fig). Other parameters including Rrs (respiratory system resistance), Rn (airway resistance), and H (tissue resistance) were unaffected by the CS or DT treatment (Fig 3E, 3F, and 3H). Collectively, these results suggest that the presence of *Arf*-expressing cells exacerbates CS-induced lung dysfunction through enhancements in the recruitment of macrophages after CS exposure.

**Fig 3.**
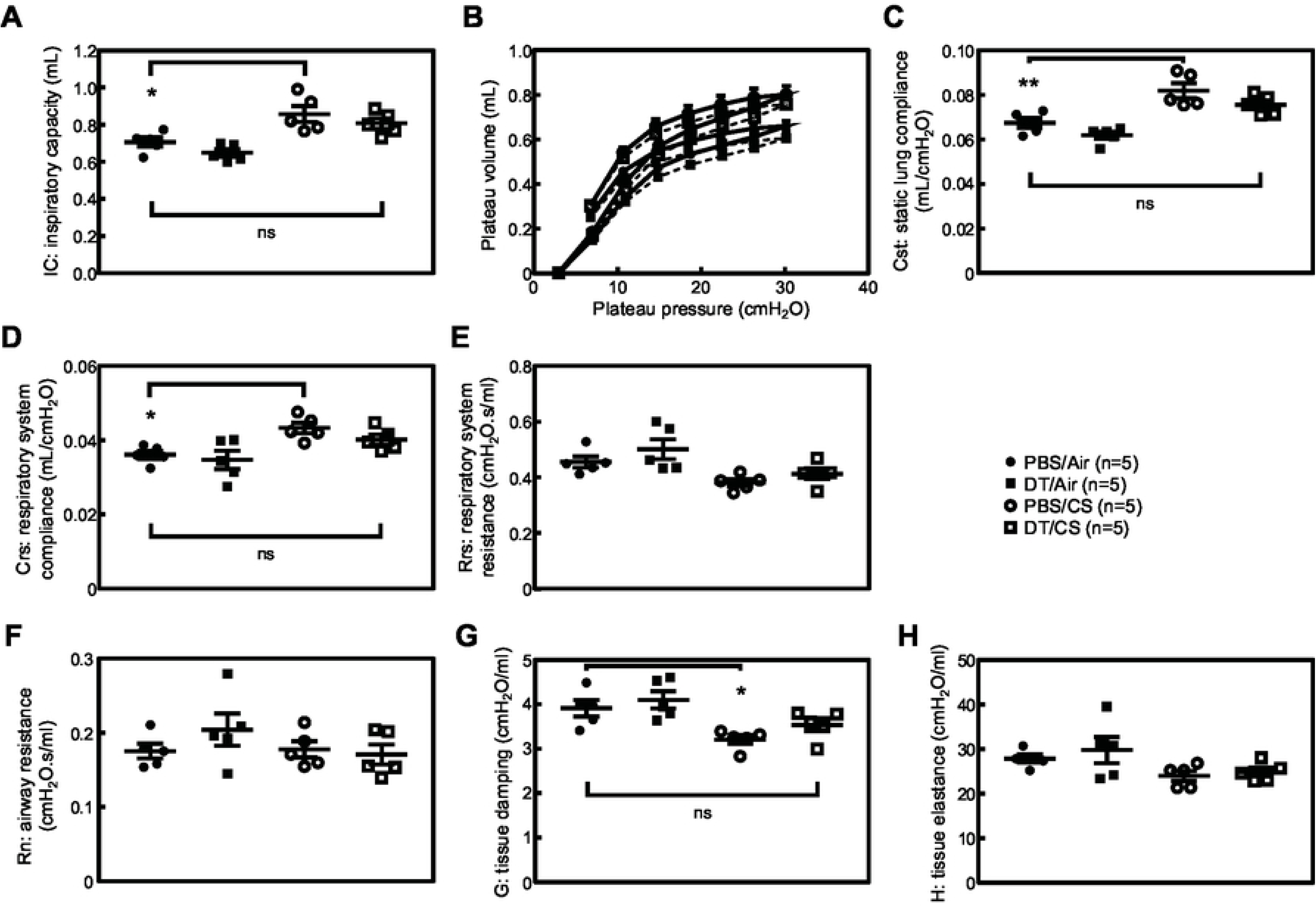
Pulmonary function was maintained in the absence of *Arf*-expressing cells after CS exposure. (**A**) Inspiratory capacity was measured in each mouse. (**B**) Pressure-volume curves of ARF-DTR mice lungs. (**C**) Static lung compliance. (**D-F**) Airway resistance (Rn), tissue damping (G), and tissue elastance (H) were analyzed. Data represent means ± SEM. Data were analyzed by a one-way ANOVA and Tukey post-hoc analysis. \**P*<0.05, \*\**P*<0.01 and ns; not significant.

### Effects of p19^Arf^-expressing cell ablation in cigarette smoke extract-treated mice

The present results and previous findings suggest that the elimination of p19^Arf^-expressing cells prior to the CS exposure or elastase inhalation exerts protective effects against emphysema-associated lung pathologies [20]. However, it currently remains unclear whether the elimination of p19^Arf^-expressing cells exerts therapeutic effects after lung injury. To test this possibility, we treated mice with DT after a damage-causing stimulation in ARF-DTR mice. In this experiment, we employed CSE inhalation as a lung injury model. CSE contains fewer chemical components than CS [30], but is sufficient to induce pulmonary inflammation [31]. Additionally, CSE has been shown to induce cellular senescence in cultured cells [32]. CSE was intranasally administered to ARF-DTR or wild-type mice for 8 weeks (3 times per week), as shown in Fig 4A. In this experiment, DT was administered 4 weeks after the CSE stimulation.

**Fig 4.**
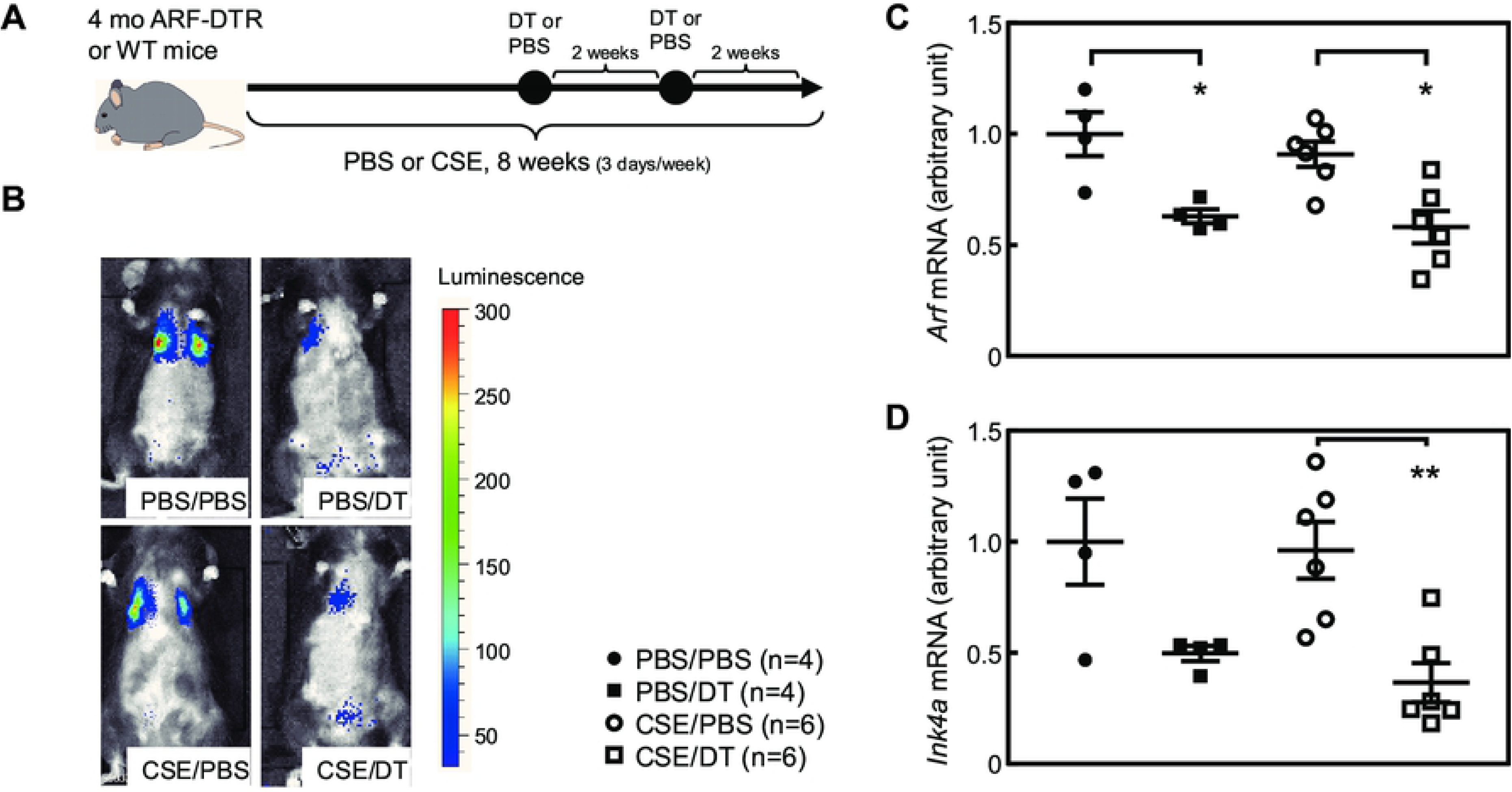
CSE inhalation model. (**A**) Experimental schedule. CSE or PBS was intranasally administered 3 times per week for 8 weeks to 4-month-old ARF-DTR or wild-type mice. DT was intraperitoneally injected twice with a 2-week interval. (**B**) Representative images of an *in vivo* imaging analysis of luciferase expression. (**C** and **D**) The expression of *Arf* (C) and *Ink4a* (D) mRNA was analyzed by real-time PCR in the lung tissue of ARF-DTR mice. Data were normalized to *Gapdh* in each group. Bars represent means ± SEM. Data were analyzed by a one-way ANOVA and Tukey post-hoc analysis. \**P*<0.05 and \*\**P*<0.01.

In contrast to the CS model (Fig 1), no marked differences were observed in lung luciferase activity between PBS- and CSE-treated animals (Fig 4B). Real-time PCR analyses consistently revealed that *Arf* and *Ink4a* expression remained unchanged in CSE-treated lung tissue (Fig 4C and 4D). Nevertheless, the DT treatment significantly reduced luciferase activity as well as *Arf* and *Ink4a* expression in both control and CSE-treated animals (Fig 4B-D), but had no effect on the expression of these genes in wild-type mice (S4 Fig).

### CSE induced pulmonary dysfunction in mice

Similar to the CS exposure, the CSE treatment also leads to impaired pulmonary function [33, 34]. We performed pulmonary function tests in CSE-treated mice. The CSE treatment resulted in increases in IC and Cst (Fig 5A-C). Other parameters including Crs, Rrs, Rn, G, and H were unaffected by the CSE treatment (Fig 5 D-H), suggesting that CSE exerts differential effects on pulmonary function or is less effective than in the CS-exposure model. IC and Cst were both restored in the DT-treated group (Fig 5A-C), while DT had no effect on these parameters in CSE-treated wild-type animals (S5 Fig).

**Fig 5.**
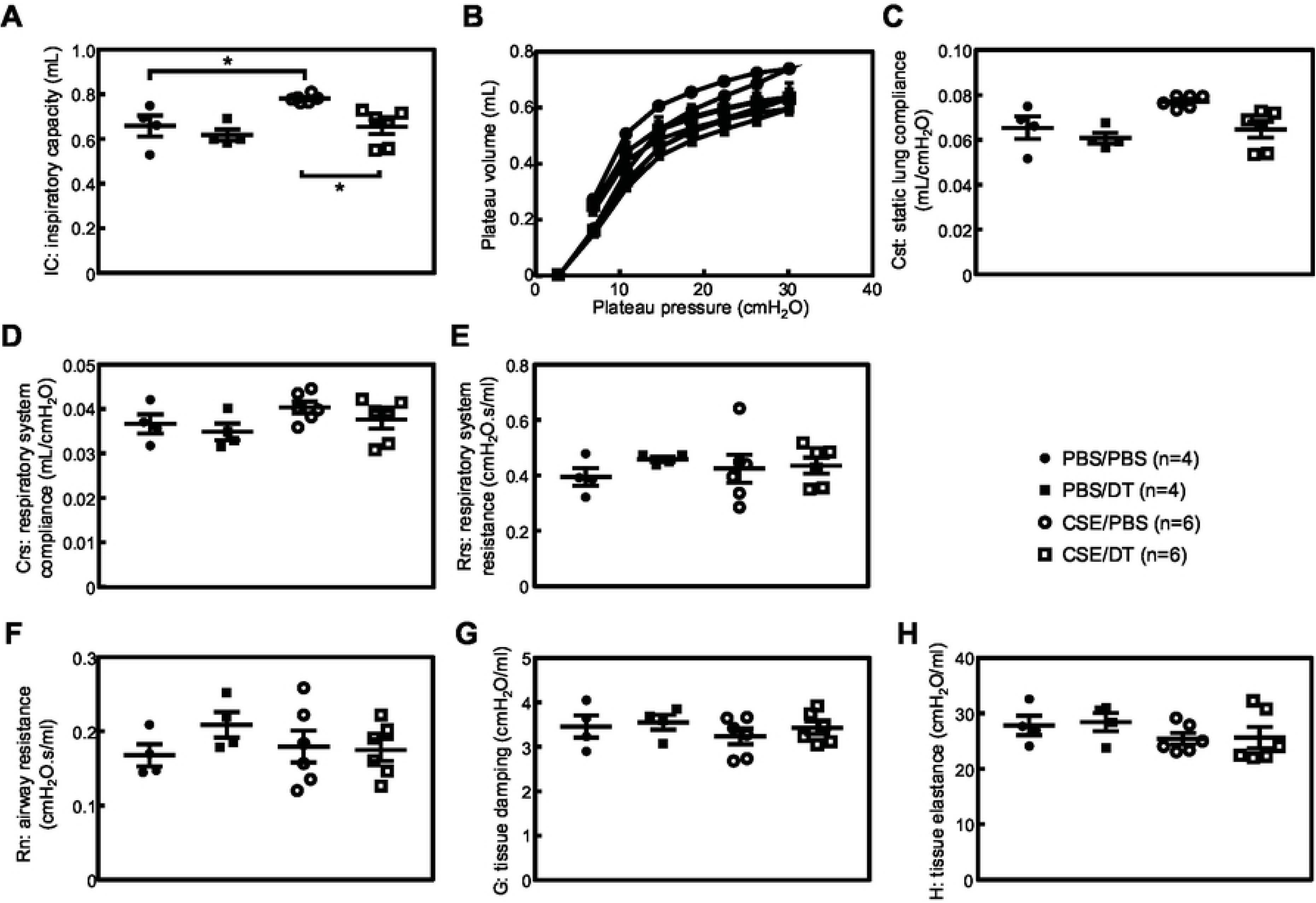
Effects of CSE on pulmonary function in ARF-DTR mice. Inspiratory capacity (A), pressure-volume loop (B), static lung compliance (C), respiratory system compliance (D), respiratory system resistance (E), airway resistance (F), tissue damping (G), and tissue elastance (H) in ARF-DTR mice were shown. Data represent means ± SEM. Data were analyzed by a one-way ANOVA and Tukey post-hoc analysis. \**P*<0.05.

We then analyzed cells in the BALF of these mice. While we and others have previously shown that the total cell number in BALF was significantly increased in the elastase-induced lung injury model [20, 28, 35], no consistent change was observed in the number of BALF cells (Fig 6A). This was somewhat expected because the continuous inhalation of PBS was sufficient to increase BALF cells [24]. Consistent with this observation, the number of macrophages, which accounts for more than 80% of the total BALF cell number, was unchanged by the CSE or DT treatment (Fig 6B). Other cells, including neutrophils, lymphocytes, and eosinophils, were also unaffected by the CSE treatment (Fig 6C-E). While the administration of DT reduced the macrophage number in CS-treated ARF-DTR mice (Fig 2) and the elastase-induced emphysema model [20], it had no effect in CSE-treated ARF-DTR mice (Fig 6B). Nevertheless, we observed a significant decrease in the neutrophil number in both PBS and CSE-treated ARF-DTR mice by DT (Fig 6D), but not in CSE-treated wild-type mice (S6 Fig). Collectively, these results suggest that CS and CSE differentially induce lung pathologies, while both stimulations lead to pulmonary dysfunction exemplified by increased lung tissue compliance, and the ablation of *Arf*-expressing cells exerts beneficial effects, even after the damage-inducing stimulation.

**Fig 6.**
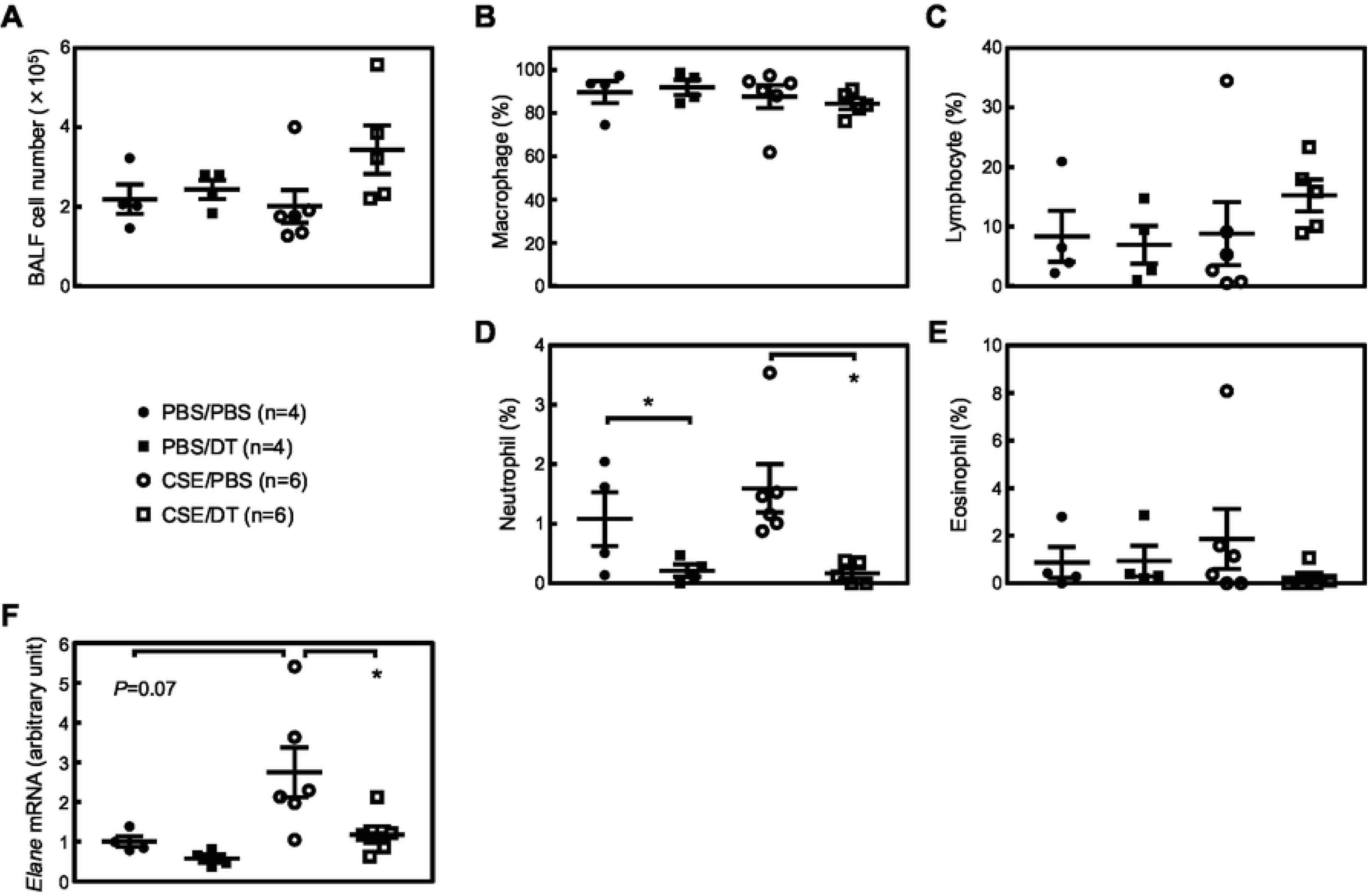
Effects of p19^Arf^ ablation on the BALF cell population. (**A**) The numbers of cells recovered from the BALF of ARF-DTR mice are shown. (**B**-**E**) The populations of macrophages (B), neutrophils (C), eosinophils (D), and lymphocytes (E) in BALF cells are shown. (**F**) The expression of neutrophil elastase (*Elane*) was analyzed by real-time PCR. Data were normalized to *Gapdh* in each sample. Data represent means ± SEM. Data were analyzed by a one-way ANOVA and Tukey post-hoc analysis. \**P*<0.05.

## Discussion

The present study demonstrated that the elimination of *Arf*-expressing cells ameliorated CS-induced lung pathologies using ARF-DTR mice. In contrast to INK-ATTAC and p16-3MR mice [36, 37], ARF-DTR mice express the transgene (DTR and luciferase) under the control of the *Arf* promoter/enhancer [19]. While the role of *ARF* in human cellular senescence and aging remains controversial, *Arf* is essential for senescence in rodent cells [38]. Additionally, the DT treatment decreased *Ink4a* as well as *Arf* expression, indicating *Arf*- and *Ink4a*-positive senescent cells were eliminated from the lung tissue of ARF-DTR mice. The elimination of *Arf*-expressing cells prior to the CS inhalation protected against CS-induced alveolar enlargement, macrophage infiltration, and pulmonary dysfunction. Additionally, the present results obtained with the CSE inhalation model suggest that the elimination of *Arf*-expressing cells after the stimulation still exerted beneficial effects on lung function.

The etiology of COPD is complex because many factors contribute to the development of this disease [1, 2] and its biology remains poorly understood. While cigarette smoking is the predominant risk factor for COPD, a relatively small proportion of smokers develop the disease. This underscores other factors in addition to CS influencing susceptibility to the disease. The incidence of COPD is increased by aging irrespective of the smoking history [39], which highlights aging-associated changes in pulmonary and/or other organs facilitating the development of this disease. Cellular senescence may be a key factor that promotes disease-associated pathologies. Cellular senescence is accelerated in emphysema patients [7], and we previously suggested that cellular senescence plays a role in CS-induced lung pathologies [29], while aging-associated changes other than cellular senescence may also contribute to the CS-induced pathology [40].

We previously reported that the elimination of *Arf*-expressing cells protected against elastase-induced emphysema [20]. Elastase inhalation is a facile and highly reproducible model for the induction of pulmonary emphysema, and a single shot of elastase is sufficient to induce alveolar collapse. However, the elastase treatment results in panlobular (panacinar) emphysema, which is a rare disease in humans. In humans, centrilobular (centriacinar) emphysema is the most common type of pulmonary emphysema, the main cause of which is smoking. Therefore, we investigated the effects of senolysis in the CS mouse model in the present study. In contrast to the elastase model, CS exposure resulted in weak, but chronic damage to the lung tissue, which leads to impaired pulmonary function. We also employed CSE inhalation for the lung injury model. CSE contains fewer chemical components than CS [30]. However, in contrast to the CS model in which the whole body is exposed to CS, the CSE inhalation model may more specifically target the tissue/organ. The present results suggest that CSE inhalation was sufficient to induce pulmonary dysfunction; however, it did not induce notable histological changes as previously documented [31]. In both models, the elimination of *Arf*-expressing cells was proven to ameliorate lung damage, suggesting that senescent cells also play an important role in CS-induced pathologies.

Inflammation in lung tissue is a critical step for the development of disease. The inactivation of macrophage or neutrophil elastase has been shown to confer resistance to CS-induced emphysema in mice [5, 6]. The results obtained herein indicate that CS and CSE differentially affect alveolar inflammatory cells. Nevertheless, the elimination of *Arf*-expressing cells suppressed the expression of elastolytic enzymes, suggesting that senescent cells exacerbate CS/CSE-induced lung pathologies by enhancing alveolar inflammation. Hall and colleagues recently reported that p16^Ink4a^ is expressed in non-senescent macrophages [41, 42]. These findings suggested that the DT treatment ablated non-senescent macrophages, thereby affecting pulmonary inflammation and the associated phenotypes. However, this is unlikely because endogenous *Arf* levels are extremely low and the expression of the transgene (DTR and luciferase) was not detected in the macrophages of ARF-DTR mice [19]. Furthermore, our previous findings indicated that p19^Arf^ was present in the mesenchymal cells of lung tissue; however, the population of *Arf*-expressing cells was estimated to be as small as 2% of all mesenchymal cells in lung tissue. Senescent cells produce chemokines/cytokines as part of SASP, which have a marked impact on the inflammatory process [43]. Hence, it is reasonable to assume that the effects of DT were attributed to the clearance of senescent cells that may facilitate the inflammatory response through SASP.

We previously demonstrated that the elimination of *Arf*-expressing cells exerted preventive effects against elastase-induced pathologies [20]. The present results obtained with the CSE model strongly suggest that senolysis also exhibits therapeutic efficacy because DT partly restored pulmonary function in ARF-DTR mice in which lung injury was induced by CSE in advance. Thus, these results suggest the potential of the application of senolytic drugs as emphysema therapy. In this regard, it is important to note that the oral administration of ABT-263/Navitoclax exerted beneficial effects against elastase-induced emphysema [20]. While ABT-263/Navitoclax have side effects [44, 45] and may not be directly applied to senolysis in humans, the targeting of senescent cells will be an effective approach. Further studies are required to assess the therapeutic efficacy of senolysis and senolytic drugs for the treatment of emphysema.

## Acknowledgments

We thank Yuko Tottori and Noboru Ogiso for their technical assistance. We also thank Dr. Mitsuo Maruyama for his financial support. This work was supported by grants from the National Center for Geriatrics and Gerontology (19-14 to MS), JSPS KAKENHI (16K08604 to MS), and the Smoking Research Foundation (to MS).

## Author contributions

R.M. performed most of the experiments. Y.S. and H.B. established the CS model. M.S. and T.S. validated the data and wrote the manuscript.

## Supporting information

**S1 Fig. *Arf* and *Ink4a* mRNA levels in CS-exposed wild-type mice.** *Arf* (A) and *Ink4a* (B) mRNA were analyzed by real-time PCR. Data were normalized to *Gapdh* mRNA in each group. Data represent means ± SEM. Data were analyzed by the Student’s *t*-test and no significance was observed.

**S2 Fig. DT had no effect on alveolar size, macrophage numbers, or *Mmp-12* expression in wild-time mice.** (**A** and **B**) Alveolar mean linear intercepts and the number of macrophages per field (×40) were measured. (**C**) The expression of *Mmp-12* was analyzed by real-time PCR. Data were normalized to *Gapdh* in each group. Data represent means ± SEM. Data were analyzed by the Student’s *t*-test and no significance was observed.

**S3 Fig.** DT had no effects on the pulmonary function of CS-exposed wild-type mice. Inspiratory capacity (A), pressure-volume loop (B), static lung compliance (C), respiratory system compliance (D), respiratory system resistance (E), airway resistance (F), tissue damping (G), and tissue elastance (H) in ARF-DTR mice were shown. Data represent means ± SEM. Data were analyzed by a one-way ANOVA and Tukey post-hoc analysis. \**P*<0.05. Data represent means ± SEM. Data were analyzed by the Student’s *t*-test and no significance was observed.

**S4 Fig.** *Arf* and *Ink4a* mRNA levels in CSE-treated wild-type mice. *Arf* (A) and *Ink4a* (B) mRNA levels were analyzed by real-time PCR. Data were normalized to *Gapdh* mRNA in each group. Data represent means ± SEM. Data were analyzed by the Student’s *t*-test and no significance was observed.

**S5 Fig.** DT had no effect on the pulmonary function of CSE-treated wild-type mice. Inspiratory capacity (A), pressure-volume loop (B), static lung compliance (C), respiratory system compliance (D), respiratory system resistance (E), airway resistance (F), tissue damping (G), and tissue elastance (H) in ARF-DTR mice were shown. Data represent means ± SEM. Data were analyzed by a one-way ANOVA and Tukey post-hoc analysis. \**P*<0.05. Data represent means ± SEM. Data were analyzed by the Student’s *t*-test and no significance was observed.

**S6 Fig.** DT had no effects on the BALF cell population in CSE-treated wild-type animals. (**A**) The numbers of cells recovered from the BALF of wild-type mice are shown. (**B**-**E**) The populations of macrophages (B), neutrophils (C), eosinophils (D), and lymphocytes (E) in BALF cells are shown. (**F**) The expression of neutrophil elastase (*Elane*) was analyzed by real-time PCR. Data were normalized to *Gapdh* in each sample. Data represent means ± SEM. Data were analyzed by the Student’s *t*-test and no significance was observed.

## References

1. Vestbo J, Hurd SS, Agusti AG, Jones PW, Vogelmeier C, Anzueto A, et al. Global strategy for the diagnosis, management, and prevention of chronic obstructive pulmonary disease: GOLD executive summary. Am J Respir Crit Care Med. 2013;187(4):347–65. doi: 10.1164/rccm.201204-0596PP. PubMed PMID: 22878278.

2. Gershon AS, Warner L, Cascagnette P, Victor JC, To T. Lifetime risk of developing chronic obstructive pulmonary disease: a longitudinal population study. Lancet. 2011;378(9795):991–6. doi: 10.1016/S0140-6736(11)60990-2. PubMed PMID: 21907862.

3. Barnes PJ, Shapiro SD, Pauwels RA. Chronic obstructive pulmonary disease: molecular and cellular mechanisms. The European respiratory journal. 2003;22(4):672–88. PubMed PMID: 14582923.

4. Barnes PJ. Mediators of chronic obstructive pulmonary disease. Pharmacol Rev. 2004;56(4):515–48. doi: 10.1124/pr.56.4.2. PubMed PMID: 15602009.

5. Hautamaki RD, Kobayashi DK, Senior RM, Shapiro SD. Requirement for macrophage elastase for cigarette smoke-induced emphysema in mice. Science. 1997;277(5334):2002–4. PubMed PMID: 9302297.

6. Shapiro SD, Goldstein NM, Houghton AM, Kobayashi DK, Kelley D, Belaaouaj A. Neutrophil elastase contributes to cigarette smoke-induced emphysema in mice. Am J Pathol. 2003;163(6):2329–35. doi: 10.1016/S0002-9440(10)63589-4. PubMed PMID: 14633606; PubMed Central PMCID: PMCPMC1892384.

7. Tsuji T, Aoshiba K, Nagai A. Alveolar cell senescence in patients with pulmonary emphysema. Am J Respir Crit Care Med. 2006;174(8):886–93. doi: 10.1164/rccm.200509-1374OC. PubMed PMID: 16888288.

8. Collado M, Blasco MA, Serrano M. Cellular senescence in cancer and aging. Cell. 2007;130(2):223–33.

9. Sherr CJ. The INK4a/ARF network in tumour suppression. NatRevMolCell Biol. 2001;2(10):731–7.

10. Ben-Porath I, Weinberg RA. The signals and pathways activating cellular senescence. Int J Biochem Cell Biol. 2005;37(5):961–76. doi: 10.1016/j.biocel.2004.10.013. PubMed PMID: 15743671.

11. Watanabe S, Kawamoto S, Ohtani N, Hara E. Impact of senescence-associated secretory phenotype and its potential as a therapeutic target for senescence-associated diseases. Cancer Sci. 2017;108(4):563–9. doi: 10.1111/cas.13184. PubMed PMID: 28165648; PubMed Central PMCID: PMCPMC5406532.

12. Dimri GP, Lee X, Basile G, Acosta M, Scott G, Roskelley C, et al. A biomarker that identifies senescent human cells in culture and in aging skin in vivo. ProcNatlAcadSciUSA. 1995;92(20):9363–7.

13. Krishnamurthy J, Torrice C, Ramsey MR, Kovalev GI, Al Regaiey K, Su L, et al. Ink4a/Arf expression is a biomarker of aging. JClinInvest. 2004;114(9):1299–307.

14. He S, Sharpless NE. Senescence in Health and Disease. Cell. 2017;169(6):1000–11. doi: 10.1016/j.cell.2017.05.015. PubMed PMID: 28575665; PubMed Central PMCID: PMCPMC5643029.

15. Baker DJ, Childs BG, Durik M, Wijers ME, Sieben CJ, Zhong J, et al. Naturally occurring p16(Ink4a)-positive cells shorten healthy lifespan. Nature. 2016;530(7589):184–9. doi: 10.1038/nature16932. PubMed PMID: 26840489; PubMed Central PMCID: PMCPMC4845101.

16. Schafer MJ, White TA, Iijima K, Haak AJ, Ligresti G, Atkinson EJ, et al. Cellular senescence mediates fibrotic pulmonary disease. Nat Commun. 2017;8:14532. doi: 10.1038/ncomms14532. PubMed PMID: 28230051.

17. Justice JN, Nambiar AM, Tchkonia T, LeBrasseur NK, Pascual R, Hashmi SK, et al. Senolytics in idiopathic pulmonary fibrosis: Results from a first-in-human, open-label, pilot study. EBioMedicine. 2019. doi: 10.1016/j.ebiom.2018.12.052. PubMed PMID: 30616998.

18. Saito M, Iwawaki T, Taya C, Yonekawa H, Noda M, Inui Y, et al. Diphtheria toxin receptor-mediated conditional and targeted cell ablation in transgenic mice. Nature biotechnology. 2001;19(8):746–50. doi: 10.1038/90795. PubMed PMID: 11479567.

19. Hashimoto M, Asai A, Kawagishi H, Mikawa R, Iwashita Y, Kanayama K, et al. Elimination of p19ARF-expressing cells enhances pulmonary function in mice. JCI Insight. 2016;1(12):e87732. doi: 10.1172/jci.insight.87732. PubMed PMID: 27699227; PubMed Central PMCID: PMCPMC5033852.

20. Mikawa R, Suzuki Y, Baskoro H, Kanayama K, Sugimoto K, Sato T, et al. Elimination of p19(ARF)-expressing cells protects against pulmonary emphysema in mice. Aging Cell. 2018;17(5):e12827. doi: 10.1111/acel.12827. PubMed PMID: 30058137; PubMed Central PMCID: PMCPMC6156494.

21. Zhu Y, Tchkonia T, Fuhrmann-Stroissnigg H, Dai HM, Ling YY, Stout MB, et al. Identification of a novel senolytic agent, navitoclax, targeting the Bcl-2 family of anti-apoptotic factors. Aging Cell. 2016;15(3):428–35. doi: 10.1111/acel.12445. PubMed PMID: 26711051; PubMed Central PMCID: PMCPMC4854923.

22. Sato T, Seyama K, Sato Y, Mori H, Souma S, Akiyoshi T, et al. Senescence marker protein-30 protects mice lungs from oxidative stress, aging, and smoking. Am J Respir Crit Care Med. 2006;174(5):530–7. doi: 10.1164/rccm.200511-1816OC. PubMed PMID: 16728709.

23. Koike K, Ishigami A, Sato Y, Hirai T, Yuan Y, Kobayashi E, et al. Vitamin C prevents cigarette smoke-induced pulmonary emphysema in mice and provides pulmonary restoration. American journal of respiratory cell and molecular biology. 2014;50(2):347–57. doi: 10.1165/rcmb.2013-0121OC. PubMed PMID: 24032444.

24. Elliott MK, Sisson JH, West WW, Wyatt TA. Differential in vivo effects of whole cigarette smoke exposure versus cigarette smoke extract on mouse ciliated tracheal epithelium. Exp Lung Res. 2006;32(3-4):99–118. doi: 10.1080/01902140600710546. PubMed PMID: 16754475; PubMed Central PMCID: PMCPMC2092449.

25. Mayhan WG, Sharpe GM. Effect of cigarette smoke extract on arteriolar dilatation in vivo. Journal of applied physiology. 1996;81(5):1996–2003. doi: 10.1152/jappl.1996.81.5.1996. PubMed PMID: 8941521.

26. Kawagishi H, Hashimoto M, Nakamura H, Tsugawa T, Watanabe A, Kontoyiannis DL, et al. HuR maintains a replicative life span by repressing the ARF tumor suppressor. MolCell Biol. 2013;33(10):1886–900.

27. Shalaby KH, Gold LG, Schuessler TF, Martin JG, Robichaud A. Combined forced oscillation and forced expiration measurements in mice for the assessment of airway hyperresponsiveness. Respiratory research. 2010;11:82. doi: 10.1186/1465-9921-11-82. PubMed PMID: 20565957; PubMed Central PMCID: PMC2904286.

28. Ueno M, Maeno T, Nishimura S, Ogata F, Masubuchi H, Hara K, et al. Alendronate inhalation ameliorates elastase-induced pulmonary emphysema in mice by induction of apoptosis of alveolar macrophages. Nat Commun. 2015;6:6332. doi: 10.1038/ncomms7332. PubMed PMID: 25757189.

29. Suzuki Y, Sato T, Sugimoto M, Baskoro H, Karasutani K, Mitsui A, et al. Hydrogen-rich pure water prevents cigarette smoke-induced pulmonary emphysema in SMP30 knockout mice. Biochem Biophys Res Commun. 2017;492(1):74–81. doi: 10.1016/j.bbrc.2017.08.035. PubMed PMID: 28807355.

30. Stedman RL. The chemical composition of tobacco and tobacco smoke. Chem Rev. 1968;68(2):153–207. PubMed PMID: 4868017.

31. Miller LM, Foster WM, Dambach DM, Doebler D, McKinnon M, Killar L, et al. A murine model of cigarette smoke-induced pulmonary inflammation using intranasally administered smoke-conditioned medium. Exp Lung Res. 2002;28(6):435–55. doi: 10.1080/01902140290096728. PubMed PMID: 12217211.

32. Nyunoya T, Monick MM, Klingelhutz A, Yarovinsky TO, Cagley JR, Hunninghake GW. Cigarette smoke induces cellular senescence. American journal of respiratory cell and molecular biology. 2006;35(6):681–8. doi: 10.1165/rcmb.2006-0169OC. PubMed PMID: 16840774; PubMed Central PMCID: PMC2643295.

33. He S, He Z, Chen Y, Ye J, Zong D, Zhang Y, et al. C-Kit/c-Kit ligand interaction of bone marrow endothelial progenitor cells is influenced in a cigarette smoke extract-induced emphysema model. Exp Lung Res. 2013;39(6):258–67. doi: 10.3109/01902148.2013.802828. PubMed PMID: 23786491.

34. He ZH, Chen P, Chen Y, He SD, Ye JR, Zhang HL, et al. Comparison between cigarette smoke-induced emphysema and cigarette smoke extract-induced emphysema. Tob Induc Dis. 2015;13(1):6. doi: 10.1186/s12971-015-0033-z. PubMed PMID: 25814921; PubMed Central PMCID: PMCPMC4374374.

35. Limjunyawong N, Craig JM, Lagasse HA, Scott AL, Mitzner W. Experimental progressive emphysema in BALB/cJ mice as a model for chronic alveolar destruction in humans. American journal of physiology Lung cellular and molecular physiology. 2015;309(7):L662–76. doi: 10.1152/ajplung.00214.2015. PubMed PMID: 26232300; PubMed Central PMCID: PMCPMC4593839.

36. Baker DJ, Wijshake T, Tchkonia T, LeBrasseur NK, Childs BG, van de Sluis B, et al. Clearance of p16Ink4a-positive senescent cells delays ageing-associated disorders. Nature. 2011;479(7372):232–6. doi: 10.1038/nature10600. PubMed PMID: 22048312; PubMed Central PMCID: PMCPMC3468323.

37. Demaria M, Ohtani N, Youssef SA, Rodier F, Toussaint W, Mitchell JR, et al. An essential role for senescent cells in optimal wound healing through secretion of PDGF-AA. Developmental cell. 2014;31(6):722–33. doi: 10.1016/j.devcel.2014.11.012. PubMed PMID: 25499914; PubMed Central PMCID: PMC4349629.

38. Kamijo T, Zindy F, Roussel MF, Quelle DE, Downing JR, Ashmun RA, et al. Tumor suppression at the mouse INK4a locus mediated by the alternative reading frame product p19ARF. Cell. 1997;91(5):649–59.

39. Terzikhan N, Verhamme KM, Hofman A, Stricker BH, Brusselle GG, Lahousse L. Prevalence and incidence of COPD in smokers and non-smokers: the Rotterdam Study. Eur J Epidemiol. 2016;31(8):785–92. doi: 10.1007/s10654-016-0132-z. PubMed PMID: 26946425; PubMed Central PMCID: PMCPMC5005388.

40. Rashid K, Sundar IK, Gerloff J, Li D, Rahman I. Lung cellular senescence is independent of aging in a mouse model of COPD/emphysema. Sci Rep. 2018;8(1):9023. doi: 10.1038/s41598-018-27209-3. PubMed PMID: 29899396; PubMed Central PMCID: PMCPMC5998122.

41. Hall BM, Balan V, Gleiberman AS, Strom E, Krasnov P, Virtuoso LP, et al. Aging of mice is associated with p16(Ink4a)- and beta-galactosidase-positive macrophage accumulation that can be induced in young mice by senescent cells. Aging (Albany NY). 2016;8(7):1294–315. doi: 10.18632/aging.100991. PubMed PMID: 27391570; PubMed Central PMCID: PMCPMC4993332.

42. Hall BM, Balan V, Gleiberman AS, Strom E, Krasnov P, Virtuoso LP, et al. p16(Ink4a) and senescence-associated beta-galactosidase can be induced in macrophages as part of a reversible response to physiological stimuli. Aging (Albany NY). 2017;9(8):1867–84. doi: 10.18632/aging.101268. PubMed PMID: 28768895; PubMed Central PMCID: PMCPMC5611982.

43. Coppe JP, Desprez PY, Krtolica A, Campisi J. The senescence-associated secretory phenotype: the dark side of tumor suppression. AnnuRevPathol. 2010;5:99–118.

44. Wilson WH, O’Connor OA, Czuczman MS, LaCasce AS, Gerecitano JF, Leonard JP, et al. Navitoclax, a targeted high-affinity inhibitor of BCL-2, in lymphoid malignancies: a phase 1 dose-escalation study of safety, pharmacokinetics, pharmacodynamics, and antitumour activity. Lancet Oncol. 2010;11(12):1149–59. doi: 10.1016/S1470-2045(10)70261-8. PubMed PMID: 21094089; PubMed Central PMCID: PMCPMC3025495.

45. Schoenwaelder SM, Jarman KE, Gardiner EE, Hua M, Qiao J, White MJ, et al. Bcl-xL-inhibitory BH3 mimetics can induce a transient thrombocytopathy that undermines the hemostatic function of platelets. Blood. 2011;118(6):1663–74. doi: 10.1182/blood-2011-04-347849. PubMed PMID: 21673344.

